# An alternative binding mode of IGHV3-53 antibodies to the SARS-CoV-2 receptor binding domain

**DOI:** 10.1101/2020.07.26.222232

**Authors:** Nicholas C. Wu, Meng Yuan, Hejun Liu, Chang-Chun D. Lee, Xueyong Zhu, Sandhya Bangaru, Jonathan L. Torres, Tom G. Caniels, Philip J.M. Brouwer, Marit J. van Gils, Rogier W. Sanders, Andrew B. Ward, Ian A. Wilson

**Affiliations:** Department of Integrative Structural and Computational Biology, The Scripps Research Institute, La Jolla, CA 92037, USA; Department of Medical Microbiology, Amsterdam UMC, University of Amsterdam, Amsterdam Infection & Immunity Institute, 1105AZ Amsterdam, the Netherlands; Department of Microbiology and Immunology, Weill Medical College of Cornell University, New York, NY 10021, USA; IAVI Neutralizing Antibody Center, The Scripps Research Institute, La Jolla, CA 92037, USA; Consortium for HIV/AIDS Vaccine Development (CHAVD), The Scripps Research Institute, La Jolla, CA 92037, USA; The Skaggs Institute for Chemical Biology, The Scripps Research Institute, La Jolla, CA, 92037, USA

## Abstract

IGHV3-53-encoded neutralizing antibodies are commonly elicited during SARS-CoV-2 infection and target the receptor-binding domain (RBD) of the spike (S) protein. Such IGHV3-53 antibodies generally have a short CDR H3 due to structural constraints in binding the RBD (mode A). However, a small subset of IGHV3-53 antibodies to the RBD contain a longer CDR H3. Crystal structures of two IGHV3-53 neutralizing antibodies here demonstrate that a longer CDR H3 can be accommodated in a different binding mode (mode B). These two classes of IGHV3-53 antibodies both target the ACE2 receptor binding site, but with very different angles of approach and molecular interactions. Overall, these findings emphasize the versatility of IGHV3-53 in this common antibody response to SARS-CoV-2, where conserved IGHV3-53 germline-encoded features can be combined with very different CDR H3 lengths and light chains for SARS-CoV-2 RBD recognition and virus neutralization.

## INTRODUCTION

Development of an effective vaccine against severe acute respiratory syndrome coronavirus 2 (SARS-CoV-2) is perhaps the most exigent health-related priority due to the ongoing COVID-19 pandemic. However, the molecular and functional understanding of the antibody response to SARS-CoV-2 infection and vaccination is still somewhat limited, but is critical for vaccine assessment and redesign. Most SARS-CoV-2 antibodies that target the receptor-binding domain (RBD) on the spike (S) protein appear to be neutralizing (Brouwer et al., 2020; Cao et al., 2020; Robbiani et al., 2020; Rogers et al., 2020; Zost et al., 2020) and the most intuitive mechanism of neutralization is that they block binding of the host receptor angiotensin-converting enzyme 2 (ACE2).

To date, several structures of antibodies that target the ACE2-binding site on RBD have been determined (Cao et al., 2020; Ju et al., 2020; Shi et al., 2020), including some that are encoded by the IGHV3-53 gene (Barnes et al., 2020; Brouwer et al., 2020; Wu et al., 2020; Yuan et al., 2020a). Our previous study demonstrated that antibodies encoded by the IGHV3-53 gene utilize germline-encoded residues to engage the ACE2-binding site on the RBD, accounting for their frequency in shared antibody responses in SARS-CoV-2 patients (Yuan et al., 2020a). Due to structural constraints in their mode of binding through interaction with the germline-encoded heavy chain complementarity determining regions (CDR) H1 and H2, a short CDR H3 (length of ≤10 amino acids, Kabat numbering) is also a molecular signature of these IGHV3-53 antibodies (Barnes et al., 2020; Yuan et al., 2020a). Nevertheless, a small subset (around 10%) of RBD-targeting IGHV3-53 antibodies have much longer CDR H3s (15 amino acids or longer) (Barnes et al., 2020; Yuan et al., 2020a). As it was not apparent how such IGHV3-53 antibodies could retain the same binding mode and fit their longer CDR H3 into a highly restricted pocket in the RBD (Yuan et al., 2020a), we aimed to resolve this conundrum.

## RESULTS

### Two RBD-targeting IGHV3-53 antibodies with different binding modes

We determined crystal structures of two IGHV3-53 neutralizing antibodies, COVA2-04 and COVA2-39 (Brouwer et al., 2020), with different CDR H3 lengths in complex with SARS-CoV-2 RBD to 2.35 and 1.72 Å resolutions, respectively (Figure 1A and Table S1). Both antibodies were derived from a convalescent donor from Amsterdam and potently neutralize SARS-CoV-2 virus (Brouwer et al., 2020). Similar to typical RBD-targeting IGHV3-53 antibodies (Barnes et al., 2020; Wu et al., 2020; Yuan et al., 2020a), COVA2-04 has a relatively short CDR H3 of 10 amino acids, whereas COVA2-39 CDR H3 is 15 amino acids (Kabat numbering, Figure S1A). COVA2-04 has only two somatic amino-acid substitutions in the heavy chain and one in the light chain, which is encoded by IGKV3-20 (Figure S1B). COVA2-39 has three somatic mutations in the heavy chain and one in the light chain, which is encoded by IGLV2-23 (Figure S1C).

**Figure 1.**
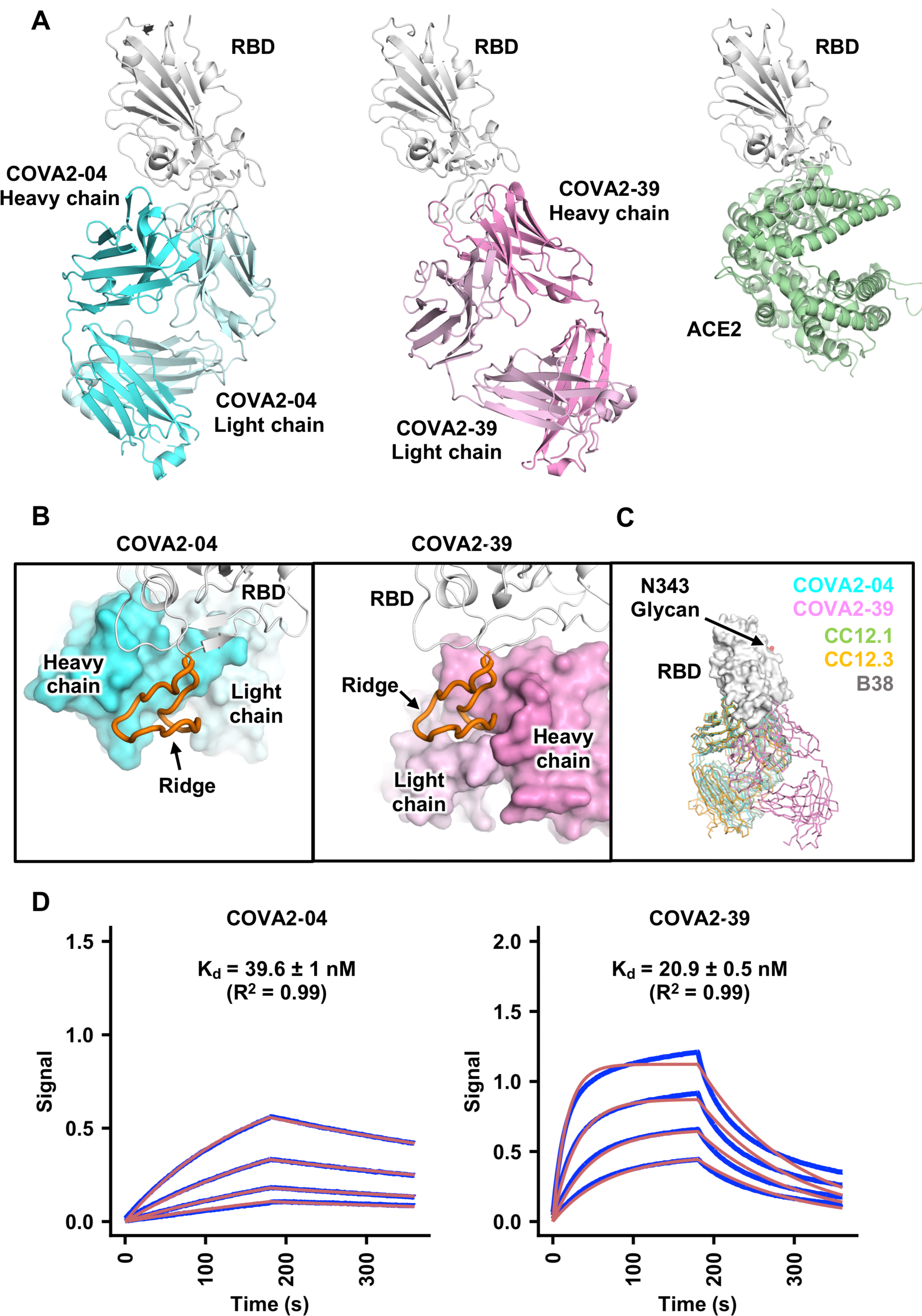
Structures of two IGHV3-53 antibodies to SARS-CoV-2 RBD with very different binding modes. **(A)** Crystal structures of COVA2-04/RBD and COVA2-39/RBD complexes are shown. Human ACE2/RBD complex is also shown for comparison (PDB 6M0J) (Lan et al., 2020). **(B)** Zoomed-in views of COVA2-04/RBD (left) and COVA2-39/RBD (right) interfaces are shown. COVA2-04 (cyan) and COVA2-39 (pink) are shown in surface representation, and RBD (white) in a cartoon representation in the same view as **A**. The ACE2-binding ridge in the RBD is in orange. **(C)** Binding modes of COVA2-04 (cyan), COVA2-39 (pink), CC12.1 (green), CC12.3 (orange), and B38 (gray) to SARS-CoV-2 (white) are compared in the same view as in **A** and **B**. CC12.1/RBD and CC12.3/RBD complexes are from PDB 6XC3 and PDB 6XC4, respectively (Yuan et al., 2020a), and RBD B38/RBD complex from PDB 7BZ5 (Wu et al., 2020). The N-glycan observed at SARS-CoV-2 RBD N343, which is distant from the epitopes of COVA2-04 and COVA2-39, is shown in red. **(E)** Binding kinetics of COVA2-04 and COVA2-39 Fabs against SARS-CoV-2 RBD were measured by biolayer interferometry (BLI). Y-axis represents the response. Blue lines represent the response curves and red lines represent a 1:1 binding model. Binding kinetics were measured for four concentrations of each Fab at 2-fold dilution starting from 125 nM. The K_d_ and R^2^ of the fitting are indicated. Representative results of two replicates are shown here.

COVA2-04 and COVA2-39 both bind to the ACE2-binding site on RBD, which is consistent with previous competition assays (Brouwer et al., 2020). Nonetheless, their angles of approach and binding modes are very different (Figure 1A-C). COVA2-04 mainly uses the light chain to interact with the flat surface of the ACE2-binding site and the heavy chain with the RBD ridge, whereas COVA2-39 mainly uses the heavy chain to interact with the flat surface and both heavy and light chains with the ridge (Figure 1B, Figure 2A-B and Table S2). In addition, COVA2-04 binds to the side of the ridge, whereas COVA2-39 binds at its tip (Figure 1B). The binding mode of COVA2-04 is very similar to previously characterized IGHV3-53 antibodies with a short CDR H3, including CC12.1, CC12.3, B38, and C105 (Barnes et al., 2020; Wu et al., 2020; Yuan et al., 2020a) (binding mode A, Figure 1C). In contrast, binding mode (mode B) of COVA2-39 is quite different and its Fab is rotated 180° along its long axis relative to COVA2-04, thereby swapping the relative orientation of the light and heavy chains, resulting in completely different molecular interactions.

**Figure 2.**
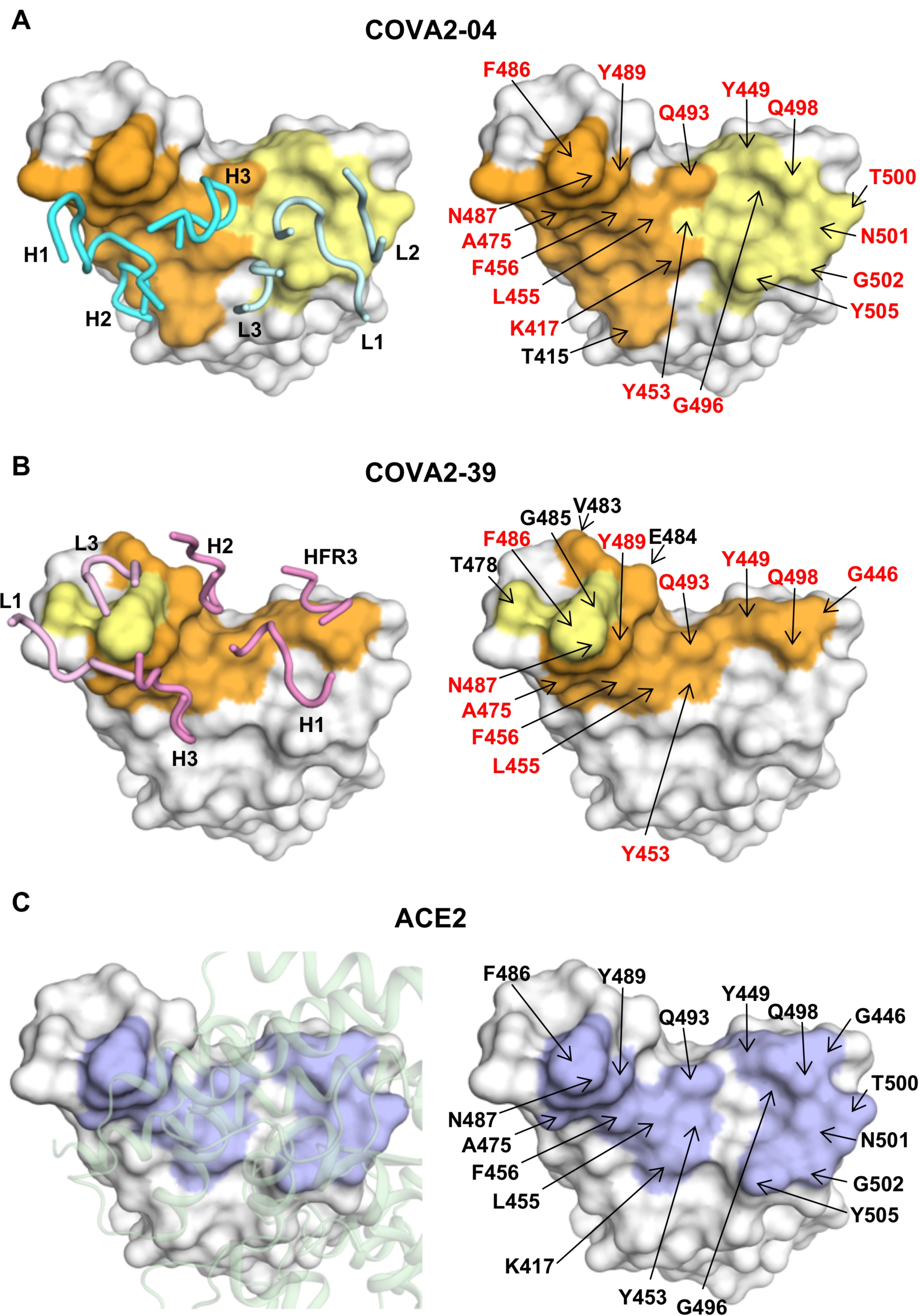
Epitopes of COVA2-04 and COVA2-39. **(A-B)** Epitope residues of **(A)** COVA2-04, and **(B)** COVA2-39 are identified by PISA (Krissinel and Henrick, 2007). Epitope residues contacting the heavy chain are in orange and light chain in yellow. On the left panels, CDR loops are labeled. On the right panels, epitope residues are labeled. For clarity, only representative epitope residues are labeled. Epitope residues that are also involved in ACE2 binding are in red. **(C)** ACE2-binding residues on the RBD are shown in blue. On the left panel, ACE2 is shown in green within a semi-transparent surface representation. On the right panel, ACE2-binding residues are labeled. A total of 17 residues in the SARS-CoV-2 RBD are used for binding by ACE2 (Lan et al., 2020). The 17 ACE2-binding residues are as described previously (Lan et al., 2020).

### Both binding modes are dominated by heavy chain

Structural modeling shows that both COVA2-04 and COVA2-39 can only bind to the RBD when it is in the “up” conformation on the trimeric S protein (Figure S2) (Walls et al., 2020; Wrapp et al., 2020). This finding is consistent with previous low-resolution, negative-stain electron microscopy analysis, which also indicated that these antibodies have different angles of approach (Brouwer et al., 2020). Despite these differences, interactions of COVA2-04 and COVA2-39 with the RBD are both dominated by the heavy chain. For COVA2-04, the buried surface areas (BSA) of the heavy and light chains are 798 Å^2^ and 360 Å^2^, respectively, compared to 576 Å^2^ and 128 Å^2^ for COVA2-39. However, this BSA difference does not translate into a corresponding difference in Fab binding affinity. Specifically, the dissociation constants (K_d_) for COVA2-04 and COVA2-39 to insect cell-expressed RBD are 40 nM and 21 nM, respectively (Figure 1D). COVA2-04 exhibits slow-on/slow-off kinetics, whereas COVA2-39 has fast-on/fast-off kinetics. Despite the faster off-rate, similar K_d_ and lower BSA, COVA2-39 IgG is more potent than COVA2-04 in neutralizing SARS-CoV-2 (IC_50_s of 0.036 and 0.22 µg/ml, respectively, in a pseudovirus assay, and 0.054 and 2.5 µg/ml in an authentic virus assay (Brouwer et al., 2020)).

Compared to COVA2-04, the COVA2-39 epitope has less overlap with the ACE2-binding site. Among 17 ACE2-binding residues on the RBD (Lan et al., 2020), 16 are within the epitope of COVA2-04 and 11 within the epitope of COVA2-39 (Figure 2A-C). The difference in angles of approach and apparent avidity of the IgG interaction in the context of the spike trimer appear to allow COVA2-39 attain higher neutralization potency, similar to avidity effects for IgG of some antibodies to influenza hemagglutinin RBD (Ekiert et al., 2012; Lee et al., 2014). Indeed, the K_d_ of IgG to mammalian cell-expressed SARS-CoV-2 spike protein is 2.3 nM for COVA2-04 and 0.1 nM for COVA2-39 (Brouwer et al., 2020), which represents a 20-fold difference in apparent IgG binding avidity that may also result from the higher local Fab concentration and rebinding of IgG to the spike protein on both the sensor in the binding experiment and on viral surface.

### Both binding modes involve similar motifs

Previously, we have described the germline-encoded features of IGHV3-53, including an _32_NY_33_ motif in CDR H1 and an _53_SGGS_56_ motif in CDR H2, which facilitate interaction with the ACE2-binding site of SARS-CoV-2 RBD in binding mode A (Yuan et al., 2020a). These motifs are also important for COVA2-04 engagement of the RBD in a similar manner to other IGHV3-53 antibodies in binding mode A (Figure 3A-B) (Yuan et al., 2020a). Interestingly, some of these germline-encoded residues are also involved in binding of COVA2-39 (mode B), but to a different location and, hence, to different residues on the RBD. V_H_ Y33 of the _32_NY_33_ motif is retained and forms a π−π stacking interaction between its aromatic ring with the aromatic side chain of Y489 (Figure 3C). Although V_H_ N32 in COVA2-39 does not interact with the RBD, both its side chain and main chain participate in a 3_10_ turn to stabilize the CDR H1 backbone (Figure 3C), similar to that observed with COVA2-04 and other IGHV3-53 antibodies (Figure 3A) (Yuan et al., 2020a). The _53_SGGS_56_ is somatically mutated to _53_TGGT_56_ in COVA2-39, which would appear to be a conservative substitution. Similar to the _53_SGGS_56_ motif in binding mode A (Figure 3B), the _53_TGGT_56_ motif in COVA2-39 also forms an extensive hydrogen-bond (H-bond) network (Figure 3D), but to a different region of SARS-CoV-2 RBD (Figure S3). The _53_TGGT_56_ motif in COVA2-39 extensively H-bonds with RBD E484 through the side chains of V_H_ T53 and V_H_ T56 (water-mediated H-bond), as well as the backbone amides of V_H_ T53, V_H_ G55, and V_H_ T56 (Figure 3D). The side chains of V_H_ T53 and V_H_ T56 also participate in additional water-mediated H-bonds with backbone carbonyls and amides of the RBD. Nevertheless, despite the similarity between Ser and Thr in size and ability to form similar H-bonds with their side-chain hydroxyl, reverting the _53_TGGT_56_ motif in COVA2-39 to the germline-encoded _53_SGGS_56_ motif decreased its K_d_ to SARS-CoV-2 RBD by at least 50-fold (Figure 1D and Figure S4A). The increased binding for _53_TGGT_56_ is likely due to the methyl groups in V_H_ T53 and T56, which make additional van der Waals interactions (Figure S4B).

**Figure 3.**
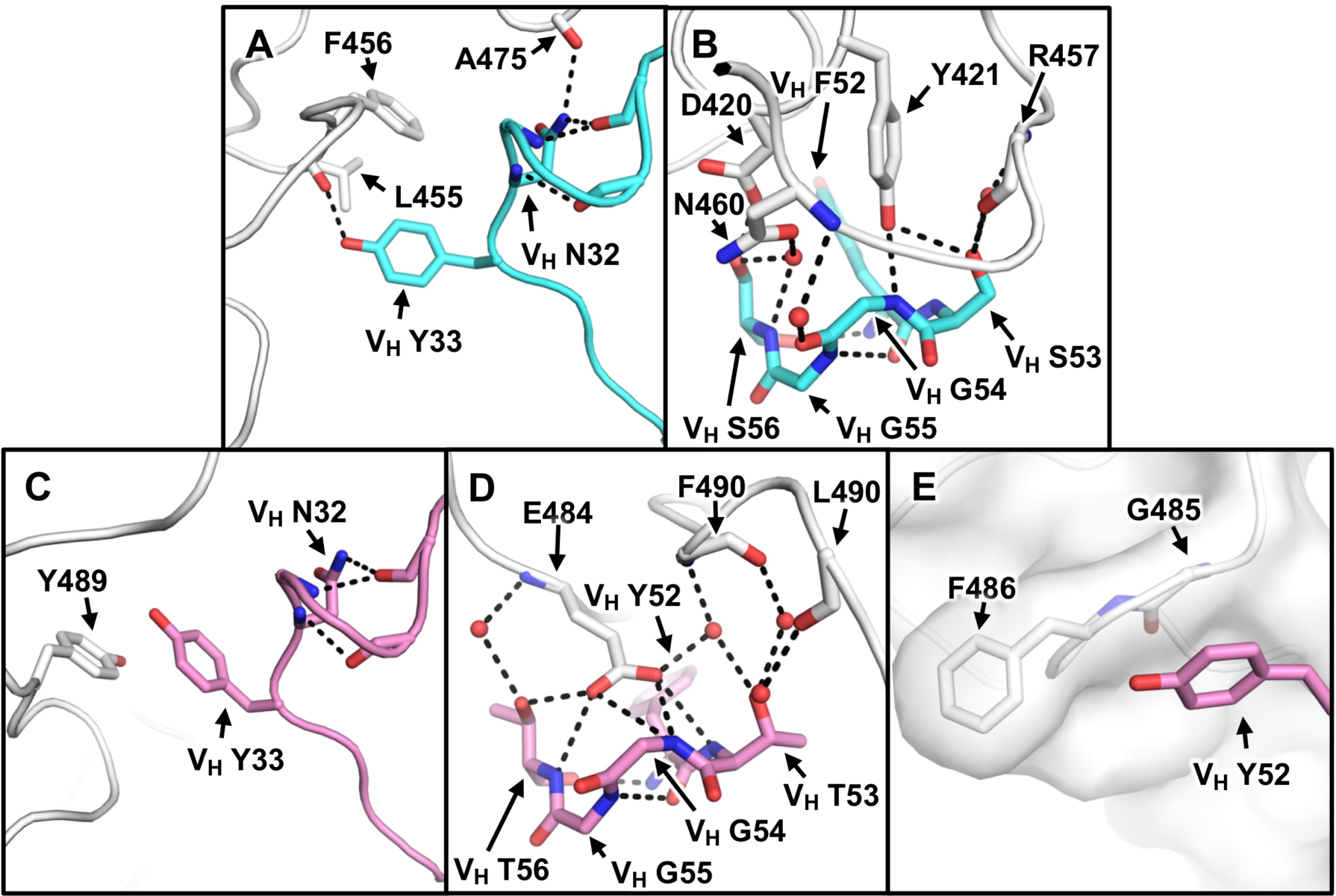
Heavy chain interactions of COVA2-39 and COVA2-04 with the RBD. **(A, C)** Interactions are shown between RBD (white) and signature _32_NY_33_ motifs on the CDR H1 loop of VH3-53 antibodies **(A)** COVA2-04 (cyan), and **(C)** COVA2-39. **(B, D)** RBD forms an extensive hydrogen bonding network with **(B)** _53_SGGS_56_ motif on the CDR H2 loop of COVA2-04, and **(D)** _53_TGGT_56_ motif on the CDR H2 loop of COVA2-39. **(F)** A π−π interaction is illustrated between G485 peptide backbone in the RBD (semi-transparent white surface) and V_H_ Y52. Hydrogen bonds are represented by dashed lines and water molecules by red spheres.

Similar to the _53_SGGS_56_ motif, _53_TGGT_56_ in COVA2-39 takes part in a type I beta turn along with V_H_ Y52, which is the first residue (i) in the turn (YTGG). The partial positive dipole from the aligned amides at the amino-end of the turn forms a charged interaction with the RBD E484 carboxyl group. Moreover, V_H_ Y52 is an important residue for binding SARS-CoV-2 RBD in binding mode B, but not in binding mode A (Figure 3E). V_H_ Y52 of COVA2-39 forms a π−π interaction with backbone peptide bond between RBD G485 and F486. COVA2-39 also employs other IGHV3-53 germline-encoded residues for interaction with the RBD, including V_H_ S30, V_H_ Y58, V_H_ N73, and V_H_ S74 (Figure S4C-D). Overall, these findings demonstrate that the germline-encoded features of IGHV3-53 are conducive for interaction with different regions of the ACE2-binding site on the RBD that involve different approach angles (i.e. binding modes A and B) in the context of different CDR H3 lengths.

### Binding mode B has less structural constraints on CDR H3 length

Next we aimed to understand the relationship between CDR H3 length and the two different binding modes. Consistent with previous structures of IGHV3-53 antibodies that target the RBD in binding mode A (Barnes et al., 2020; Wu et al., 2020; Yuan et al., 2020a), CDR H3 of COVA2-04 is highly buried by the RBD and the light chain (Figure 4A). This observation substantiates the notion that IGHV3-53 in binding mode A has strong structural constraints on CDR H3 length. In contrast, in binding mode B, the longer CDR H3 of COVA2-39 is largely solvent exposed (Figure 4B). This observation explains why IGHV3-53 can accommodate a longer CDR H3 in binding mode B (COVA2-39) compared to mode A (COVA2-04, CC12.1, CC12.3, B38, and C105). Interestingly, COVA2-39 CDR H3 interacts with the RBD mostly through non-specific van der Waals interactions. Three H-bonds are also made, two of which (one water-mediated) involve the CDR H3 backbone, whereas only one involves a side chain (V_H_ E100c) (Figure 4C). Thus, interaction between CDR H3 of COVA2-39 and RBD is largely sequence non-specific and simply accommodates its longer length. The CDR H3 sequences of IGHV3-53 antibodies in binding mode A (COVA2-04, CC12.1, CC12.3, and B38) are also quite different (Wu et al., 2020; Yuan et al., 2020a) (Figure 4D). As a result, while both binding modes A and B do not have strong constraints on the actual CDR H3 sequences, structural constraints on CDR H3 length is much stronger in binding mode A compared to mode B.

**Figure 4.**
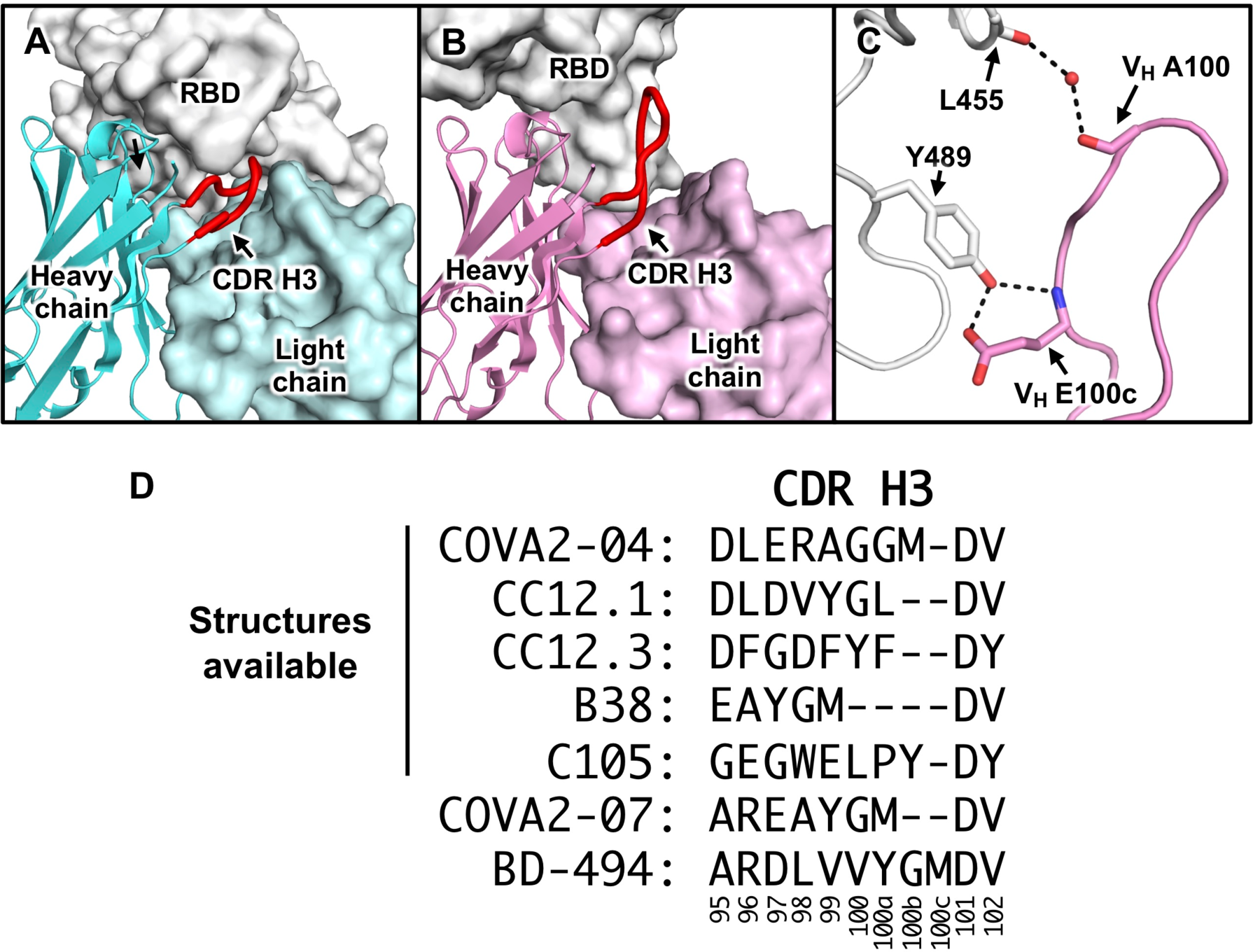
Structural constraints of CDR H3 length in different binding modes of IGHV3-53 antibodies. **(A)** Interaction between COVA2-04 (cyan) and the RBD (white) is shown with CDR H3 highlighted in red. **(B)** Interaction between COVA2-39 (pink) and the RBD (white) is shown with CDR H3 highlighted in red. In this view, the RBD is rotated ∼180° relative to (**A**). **(C)** CDR H3 of COVA2-39 makes very few contacts with the RBD. Hydrogen bond interactions are represented by dashed lines and water molecules by red spheres. **(D)** CDR H3 sequences are compared for COVA2-04, CC12.1 (Yuan et al., 2020a), CC12.3 (Yuan et al., 2020a), B38 (Wu et al., 2020), C105 (Barnes et al., 2020), COVA2-07 (Brouwer et al., 2020), and BD-494 (Cao et al., 2020). Of note, BD-494 belongs to a cluster of RBD-targeting IGHV3-53 antibodies with similar CDR H3 sequences, including BD500, BD-503, BD-505, BD-506, BD-507, and BD-508 (Cao et al., 2020). Residue positions are labeled according to the Kabat numbering scheme.

### Light chain of IGHV3-53 antibody is associated with CDR H3 length

An important feature in COVA2-39 interaction with the RBD is its engagement of the ACE2-binding ridge. Specifically, RBD F486 at the tip of the ridge is anchored in a pocket formed by both heavy and light chains (Figure 5A). RBD F486 forms a π−π stacking interaction with V_L_ Y91, which in turn is stabilized by a network of T-shaped π−π stacking interactions involving V_H_ W47, V_H_ F100g, V_L_ Y30, and V_L_ W96 (Figure 5B, Figure S5). This critical interaction with the ridge indicates that the light-chain identity is important for binding mode B. In fact, the CDR H3 lengths of RBD-targeting IGHV3-53 antibodies seem to depend on the light chain identity. For example, all known RBD-targeting IGHV3-53 antibodies that pair with IGKV1-9 (n = 20) and IGKV3-20 (n = 15) have a relatively short CDR H3 (7 to 11 amino acids) (Figure 5C) (Brouwer et al., 2020; Cao et al., 2020; Ju et al., 2020; Robbiani et al., 2020; Rogers et al., 2020; Wu et al., 2020). In contrast, all three known RBD-targeting IGHV3-53 antibodies that pair with IGLV2-23, including COVA2-39, have a CDR H3 of 14 amino acids or longer (Figure 5C). Therefore, despite the limited sample size, there seems to be a relationship between the light chain identity of RBD-targeting IGHV3-53 antibodies and CDR H3 length. This observation further supports a role for light-chain identity, together with CDR H3 length, in determining the particular binding mode of IGHV3-53 antibodies to SARS-CoV-2 RBD.

**Figure 5.**
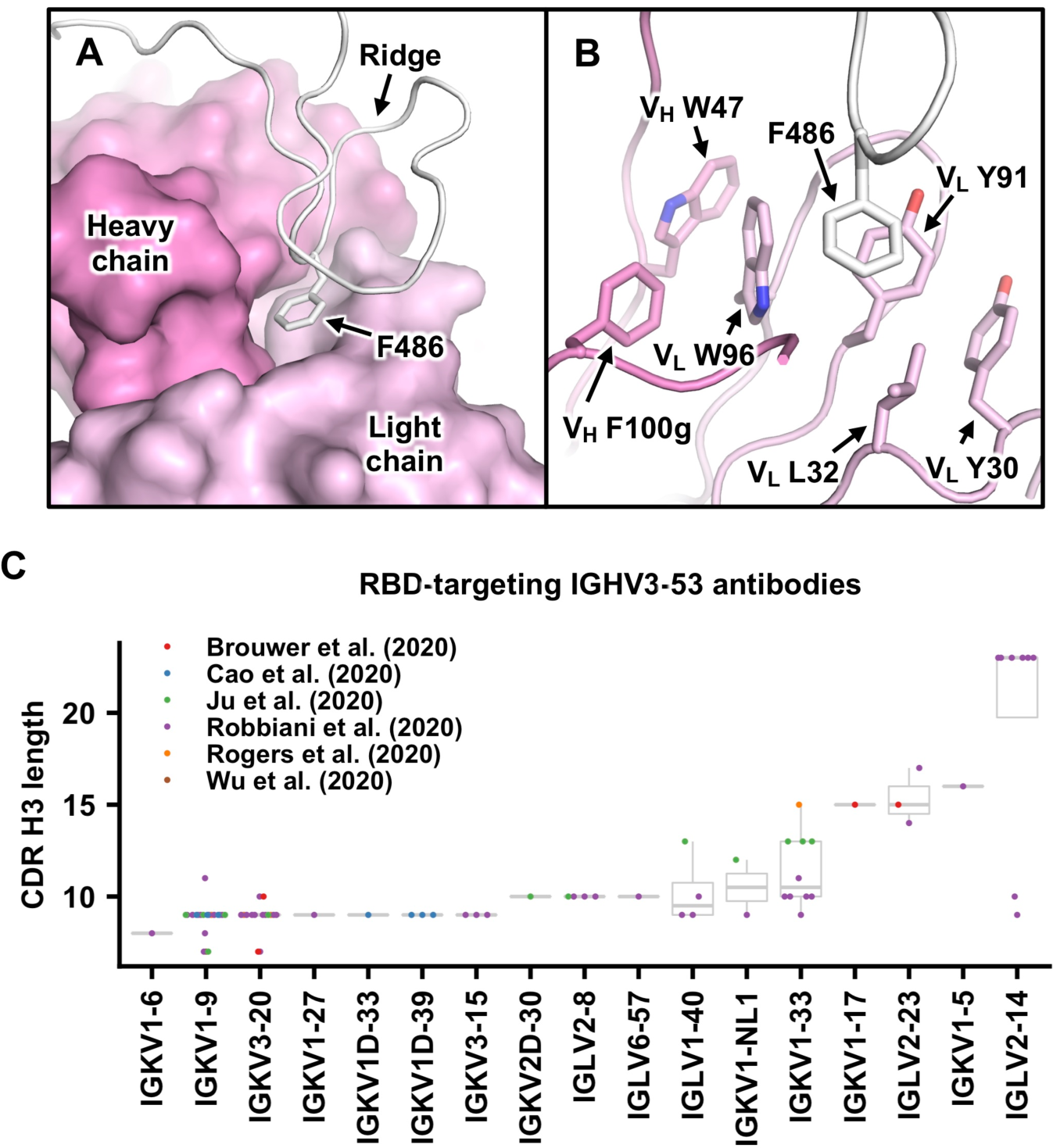
Structural analysis of the RBD ridge-anchoring pocket in COVA2-39. **(A)** The ACE2-binding ridge of the RBD (white) is shown in a cartoon representation. F486 at the tip of the ridge is shown as a stick representation and COVA2-39 Fab in a surface representation. **(B)** Interaction between F486 and COVA2-39 in the ridge-anchoring pocket is shown. **(C)** CDR H3 length of previously discovered RBD-targeting IGHV3-53 antibodies is summarized (Brouwer et al., 2020; Cao et al., 2020; Ju et al., 2020; Rogers et al., 2020; Wu et al., 2020). The light chain genes for these antibodies are shown in the bottom panel.

## DISCUSSION

The additional binding mode identified here demonstrates that IGHV3-53 is even more versatile than previously thought in SARS-CoV-2 RBD-targeting antibodies (Yuan et al., 2020a). However, a remaining question is why binding mode A is more commonly seen given the structural constraints for below average CDR H3 length, even for VH3-53 antibodies in general (Barnes et al., 2020; Wu et al., 2020; Yuan et al., 2020a). In mode A, the germline sequences of CDR H1 and H2, along with a conserved residue in FR3, seems to be the major determinant of binding without much involvement of specific residues of CDR H3. In binding mode B, formation of the RBD ridge-anchoring pocket involves both CDR H3 and L3 loops. In COVA2-39, the RBD ridge-anchoring pocket is formed from V_H_ F100g in CDR H3 and V_L_ W96 in CDR L3 and generated during V(D)J recombination. Thus, the probability of satisfying the molecular requirements to create the RBD ridge-anchoring pocket (binding mode B) could be less than simply using the unmutated germline sequence with no particular CDR H3 sequence requirements other than length (binding mode A). Nonetheless, additional structures of RBD-targeting IGHV3-53 antibodies with longer CDR H3s should further aid in explaining the differential occurrence frequency between binding modes A and B.

Given the large number of neutralizing antibodies currently being identified (Andreano et al., 2020; Brouwer et al., 2020; Cao et al., 2020; Chi et al., 2020; Ju et al., 2020; Kreer et al., 2020; Robbiani et al., 2020; Rogers et al., 2020; Seydoux et al., 2020; Zost et al., 2020), structural understanding of the antigenicity and immunogenicity of SARS-CoV-2 S protein is still at an early stage. It is clear that antibodies are elicited to the RBD in natural infection. As SARS-CoV-2 may eventually become endemic within the human population (Li et al., 2020) and escape mutations may arise, the structural information elucidated here can be harnessed for modifying or improving existing vaccine designs, and for assessing the quality and efficacy of vaccine responses.

## ACKNOWLEDGEMENTS

We thank Henry Tien for technical support with the crystallization robot, Jeanne Matteson and Yuanzi Hua for contribution to mammalian cell culture, Wenli Yu to insect cell culture, and Robyn Stanfield for assistance in data collection. We are grateful to the staff of Stanford Synchrotron Radiation Laboratory (SSRL) Beamline 12-1 for assistance. This work was supported by NIH K99 AI139445 (N.C.W.), the Bill and Melinda Gates Foundation OPP1170236 (A.B.W., I.A.W.), OPP1111923, OPP1132237, and INV-002022 (R.W.S.), and NIH HIVRAD P01 AI110657 (R.W.S., A.B.W., I.A.W.). M.J.v.G. is a recipient of an AMC Fellowship, and R.W.S is a recipient of a Vici grant from the Netherlands Organization for Scientific Research (NWO). Use of the SSRL, SLAC National Accelerator Laboratory, is supported by the U.S. Department of Energy, Office of Science, Office of Basic Energy Sciences under Contract No. DE-AC02–76SF00515. The SSRL Structural Molecular Biology Program is supported by the DOE Office of Biological and Environmental Research, and by the National Institutes of Health, National Institute of General Medical Sciences (including P41GM103393).

## AUTHOR CONTRIBUTIONS

N.C.W., M.Y., H.L. and I.A.W. conceived and designed the study. N.C.W., M.Y., H.L. and C.C.D.L. expressed and purified the proteins. T.G.C., P.J.M.B., M.J.v.G. and R.W.S. provided antibody clones and sequences. S.B., J.L.T., and A.B.W provided the nsEM maps and performed fitting. N.C.W., M.Y., H.L. and X.Z. performed the crystallization, X-ray data collection, determined and refined the X-ray structures. N.C.W., M.Y., H.L., C.C.D.L., X.Z. and I.A.W. analyzed the data. N.C.W., M.Y., H.L. and I.A.W. wrote the paper and all authors reviewed and/or edited the paper.

## MATERIALS AND METHODS

### Expression and purification of SARS-CoV-2 RBD

The receptor-binding domain (RBD) (residues 319-541) of the SARS-CoV-2 spike (S) protein (GenBank: QHD43416.1) was cloned into a customized pFastBac vector (Ekiert et al., 2011), and fused with an N-terminal gp67 signal peptide and C-terminal His_6_ tag (Yuan et al., 2020b). A recombinant bacmid DNA was generated using the Bac-to-Bac system (Life Technologies). Baculovirus was generated by transfecting purified bacmid DNA into Sf9 cells using FuGENE HD (Promega), and subsequently used to infect suspension cultures of High Five cells (Life Technologies) at an MOI of 5 to 10. Infected High Five cells were incubated at 28 °C with shaking at 110 r.p.m. for 72 h for protein expression. The supernatant was then concentrated using a 10 kDa MW cutoff Centramate cassette (Pall Corporation). The RBD protein was purified by Ni-NTA, followed by size exclusion chromatography, and buffer exchanged into 20 mM Tris-HCl pH 7.4 and 150 mM NaCl.

### Expression and purification of Fabs

For COVA2-04 and COVA2-39, the heavy and light chains were cloned into phCMV3. The plasmids were transiently co-transfected into ExpiCHO cells at a ratio of 2:1 (HC:LC) using ExpiFectamine™ CHO Reagent (Thermo Fisher Scientific) according to the manufacturer’s instructions. The supernatant was collected at 10 days post-transfection. The Fabs were purified with a CaptureSelect™ CH1-XL Affinity Matrix (Thermo Fisher Scientific) followed by size exclusion chromatography.

### Crystallization and structural determination

COVA2-04/RBD and COVA2-39/RBD complexes were formed by mixing each of the protein components at an equimolar ratio and incubated overnight at 4°C. Each complex was adjusted to 12 mg/ml and screened for crystallization using the 384 conditions of the JCSG Core Suite (Qiagen) on our custom-designed robotic CrystalMation system (Rigaku) at Scripps Research. Crystallization trials were set-up by the vapor diffusion method in sitting drops containing 0.1 μl of protein and 0.1 μl of reservoir solution. Diffraction-quality crystals were obtained in the following conditions:

COVA2-04/RBD complex (12 mg/mL): 8.5% isopropanol, 10% ethylene glycol, 15% glycerol, 0.085 M HEPES pH 7.5, and 17% polyethylene glycol 4000 at 20 °C. COVA2-39/RBD complex (12 mg/mL): 0.1 M sodium citrate pH 5.6, 20% isopropanol, 10% ethylene glycol, and 20% polyethylene glycol 4000 at 20 °C.

All crystals appeared on day 3, harvested on day 7, and were then flash cooled and stored in liquid nitrogen until data collection. Diffraction data were collected at cryogenic temperature (100 K) at Stanford Synchrotron Radiation Lightsource (SSRL) on the new Scripps/Stanford beamline 12-1 with a beam wavelength of 0.97946 Å, and processed with HKL2000 (Otwinowski and Minor, 1997). Structures were solved by molecular replacement using PHASER (McCoy et al., 2007). The models for molecular replacement of RBD and COVA2-04 were from PBD 6XC4 (Yuan et al., 2020a), whereas the model of COVA2-39 was generated by Repertoire Builder (https://sysimm.ifrec.osaka-u.ac.jp/rep_builder/) (Schritt et al., 2019). Iterative model building and refinement were carried out in COOT (Emsley et al., 2010) and PHENIX (Adams et al., 2010), respectively. Epitope and paratope residues, as well as their interactions, were identified by accessing PISA (Proteins, Interfaces, Structures and Assemblies) at the European Bioinformatics Institute (http://www.ebi.ac.uk/pdbe/prot_int/pistart.html) (Krissinel and Henrick, 2007).

### Biolayer interferometry binding assay

Antibody binding and competition assays were performed by biolayer interferometry (BLI) using an Octet Red instrument (FortéBio) as described previously (Wu et al., 2017), with anti-human Fab-CH1 2nd generation (FAB2G) biosensors. There were five steps in the assay: 1) baseline: 60 s with 1x kinetics buffer; 2) loading: 240 s with 50 μg/mL of COVA2-04 Fab or COVA2-39 Fab; 3) baseline: 60 s with 1x kinetics buffer; 4) association: 180 s with serial diluted concentrations of SARS-CoV-2 RBD; and 5) dissociation: 180 s with 1x kinetics buffer. For K_d_ estimation, a 1:1 binding model was used.

### Data availability

X-ray coordinates and structure factors are being deposited in the RCSB Protein Data Bank.

---

**Supplementary Figure 1.**
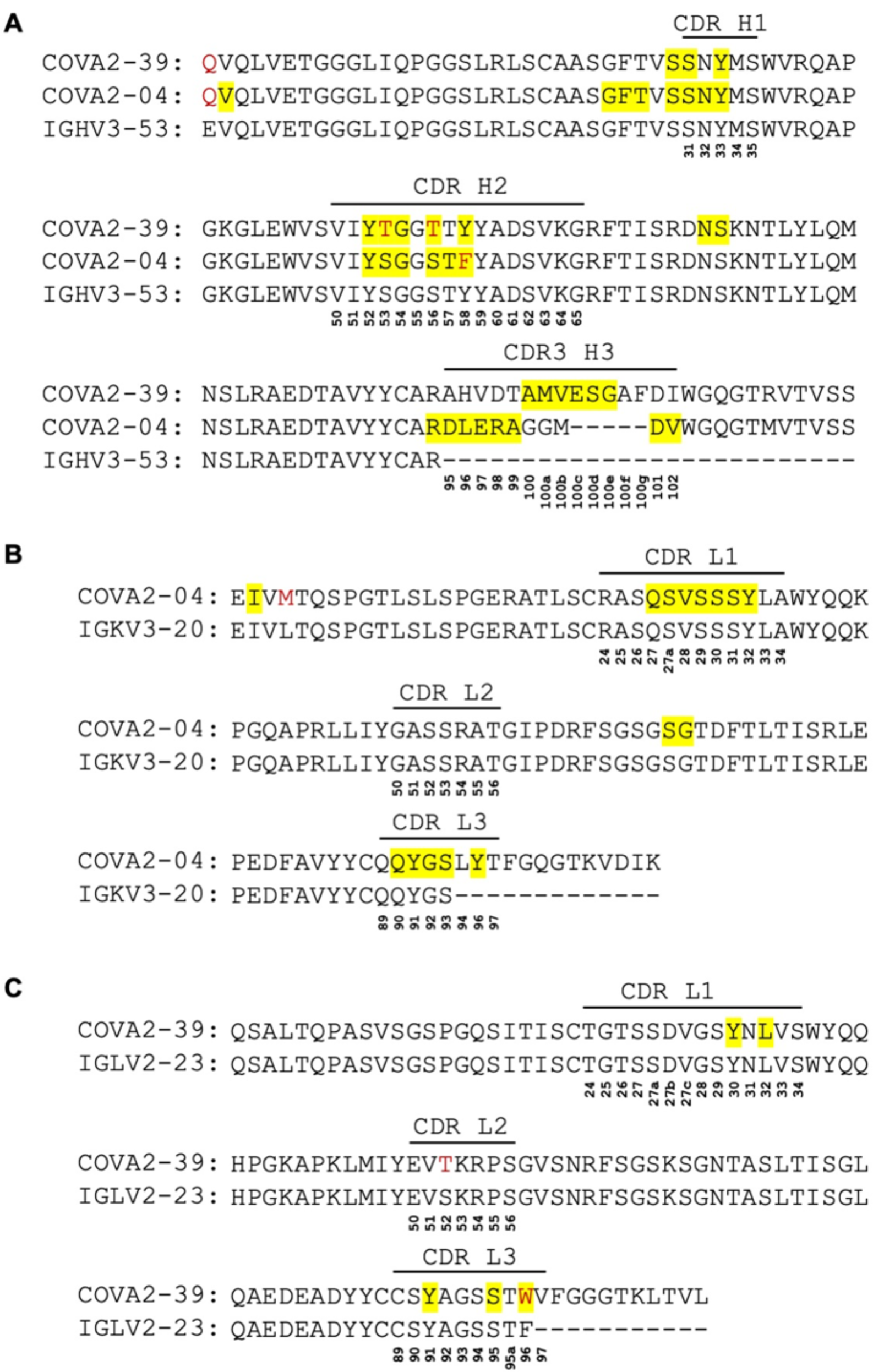
Comparison of COVA2-04 and COVA2-39 sequences to germline sequences. **(A)** Alignment of the heavy-chain variable domain sequences of COVA2-04 and COVA2-39 with the germline IGHV3-53 sequence **(B)** Alignment of the light-chain variable domain sequence of COVA2-04 with the germline IGKV3-20 sequence. **(C)** Alignment of the light-chain variable domain sequence of COVA2-39 with the germline IGLV2-23 sequence. The regions that correspond to CDR H1, H2, H3, L1, L2, and L3 are indicated. Residues that differ from the germline are highlighted in red. Residue positions in the CDRs are labeled according to the Kabat numbering scheme. Residues that interact with the RBD are highlighted in yellow.

**Supplementary Figure 2.**
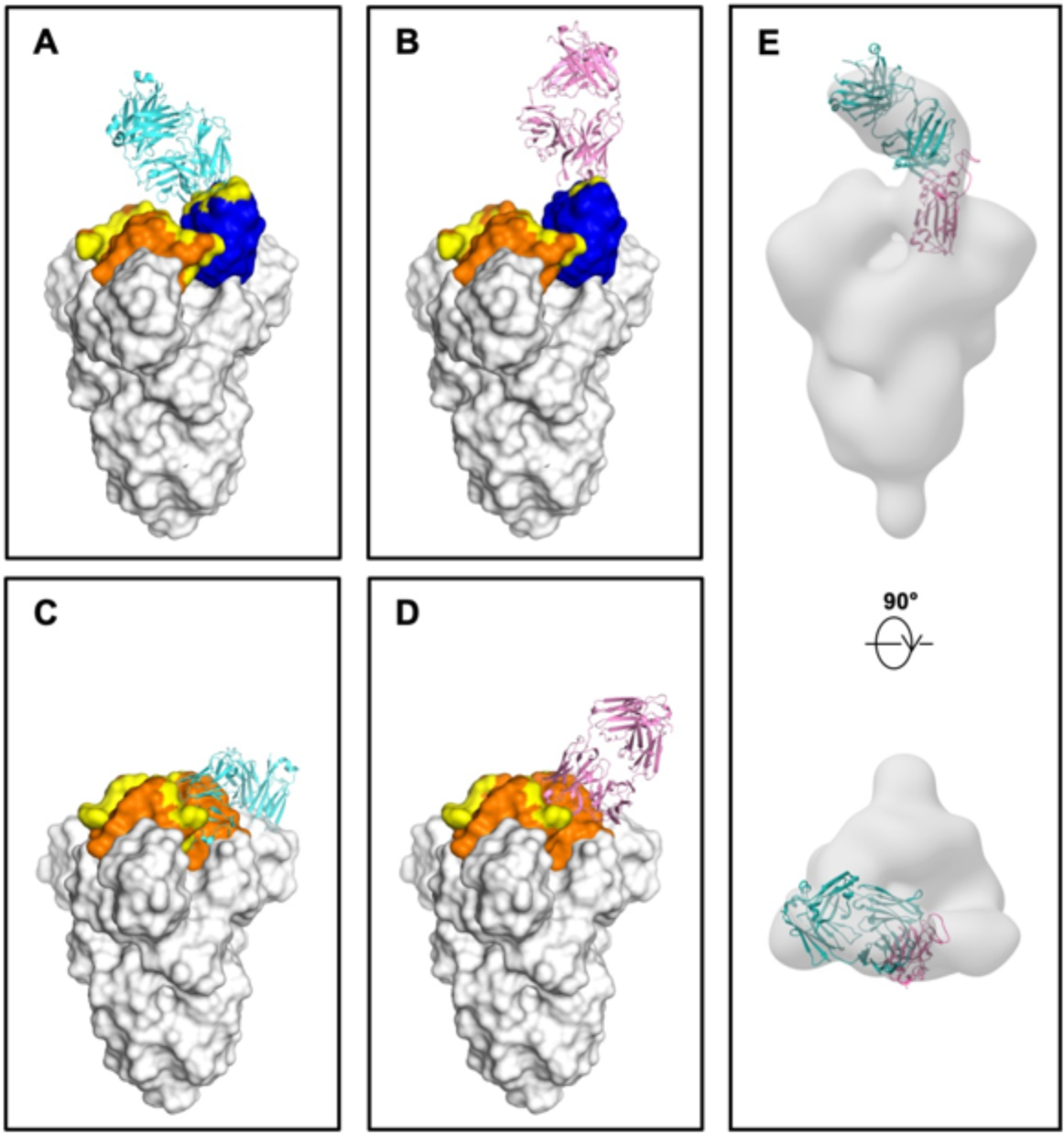
Modelling the binding of COVA2-39 and COVA2-04 on the homotrimeric SARS CoV-2 spike (S) protein. **(A-B)** The SARS-CoV-2 S trimer is shown with one RBD in the up conformation (cyan) and two RBDs in the down conformation (orange) (PDB: 6VSB) (Wrapp et al., 2020). Binding of **(A)** COVA2-04 (cyan) and **(B)** COVA2-39 (pink) to the RBDs in the up conformations is modelled. **(C-D)** The SARS-CoV-2 S trimer is shown with all three RBD in down conformations (PDB 6VXX) (Walls et al., 2020). **(C)** COVA2-04 (cyan) and **(D)** COVA2-39 (pink) cannot bind to their epitopes in the RBD in the down conformation as the epitopes are partially buried. Epitopes are colored in yellow. **(E)** COVA2-04/RBD crystal structure was fitted here to the negative-stain electron microscopy (nsEM) reconstruction of the Fab COVA-2-04/SARS CoV-2 spike protein complex that was previously generated (Brouwer et al., 2020). A similar analysis was not performed with the COVA2-39/RBD because of the poorer quality of nsEM map of the COVA2-39/S protein complex (Brouwer et al., 2020).

**Supplementary Figure 3.**
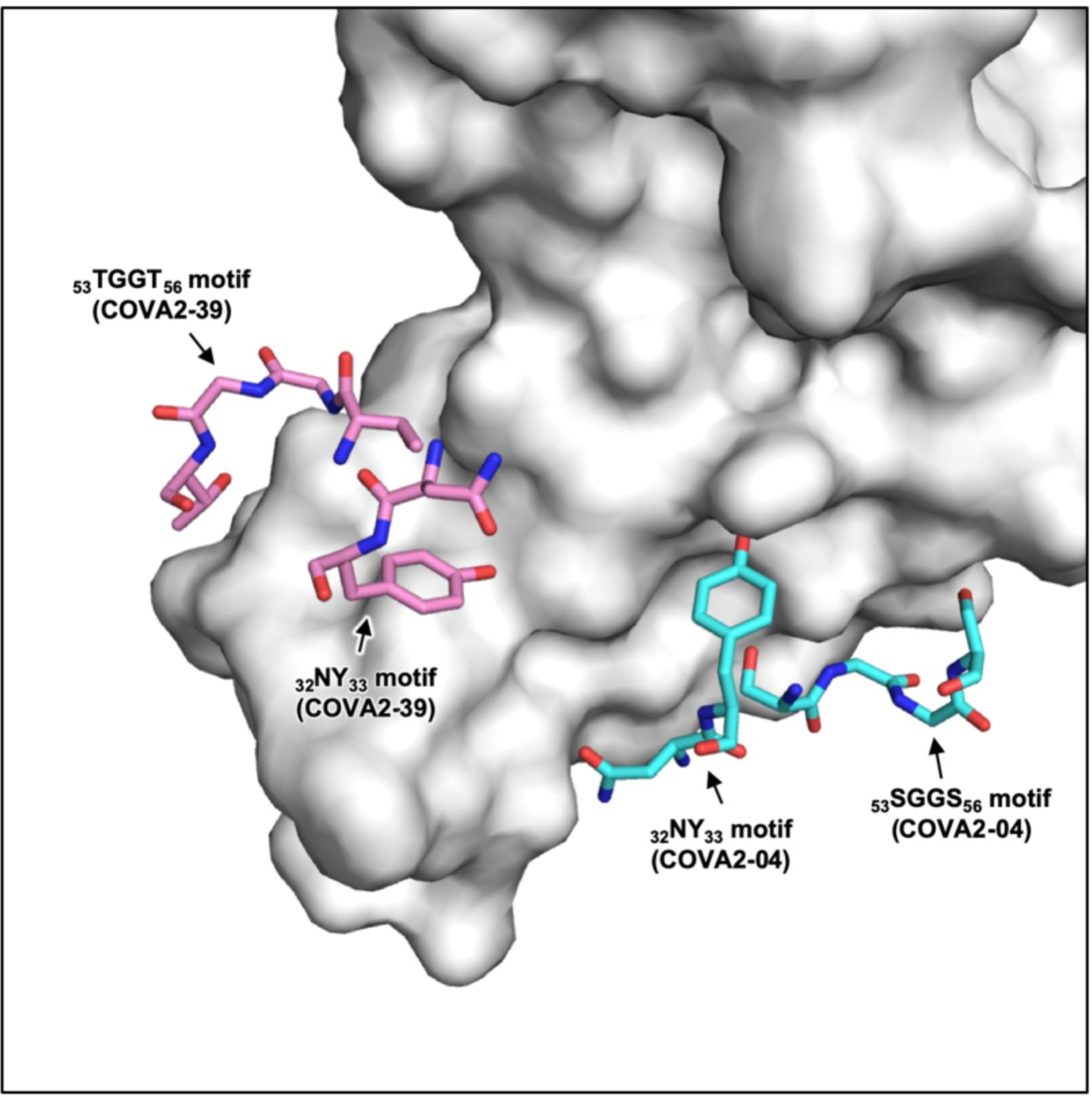
Locations of _32_NY_33_ motif and _53_SGGS_56_ (_53_TGGT_56_) motifs when VH3-53 antibodies bind the SARS CoV-2 RBD. The locations of _32_NY_33_ motif and _53_SGGS_56_ motif in COVA2-04 (cyan) as well as _32_NY_33_ motif and _53_TGGT_56_ motif in COVA2-39 (pink) are shown. SARS-CoV-2 RBD is shown as a white surface.

**Supplementary Figure 4.**
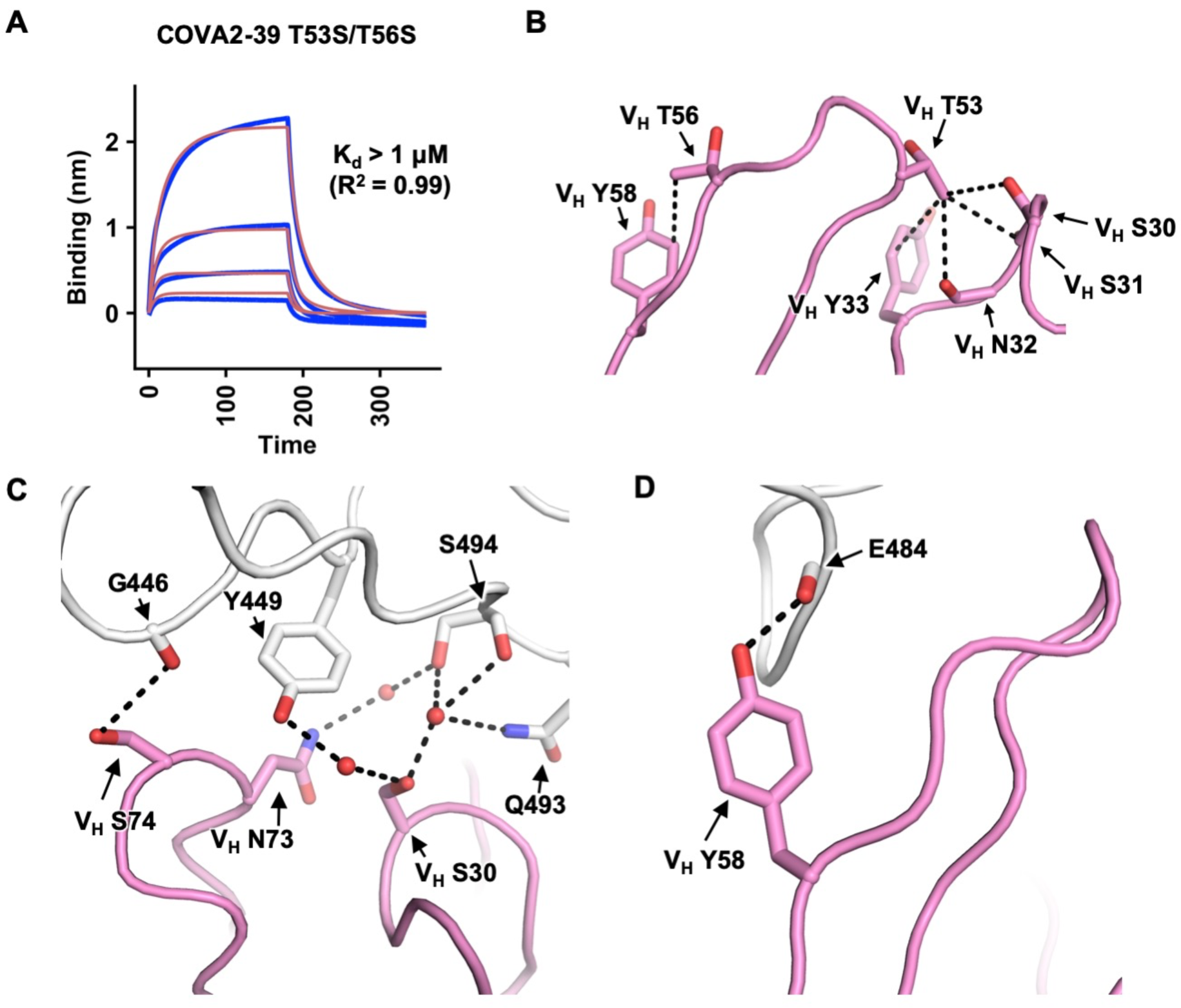
Key interactions between COVA2-39 and RBD. **(A)** Binding kinetics of COVA2-39 T53S/T56S somatic revertant in Fab format to SARS-CoV-2 RBD were measured by biolayer interferometry (BLI). Y-axis represents the response and blue lines represent the response curves. A 1:1 binding model did not fit very well, potentially due to some contribution of non-specific binding to the response curve. Subsequently, a 2:1 heterogeneous ligand model was used to improve the fit, which is represented by the red lines. In both models, the K_d_ estimated is > 1 μM. Binding kinetics were measured for four concentrations of each Fab at 2-fold dilution starting from 500 nM. The K_d_ and R^2^ of the fitting are indicated. Representative result of two replicates is shown here. **(B)** Van der Waals Interactions (within a distance of 4 Å, dashed lines) that involve the methyl group of V_H_ T53 and V_H_ T56 of COVA2-39 are shown. **(C)** The hydroxyl side chain of V_H_ S30 interacts with RBD Y449, Q493, and S494 through water-mediated H-bonds. The side chain of V_H_ N73 interacts with RBD S494 through a water-mediated H-bond. The side chain of V_H_ S74 H-bonds with the backbone carbonyl of RBD G446. **(D)** The side chain of V_H_ Y58 H-bonds with the backbone carbonyl of RBD E484. Hydrogen bonds are represented by dashed lines and water molecules by red spheres.

**Supplementary Figure 5.**
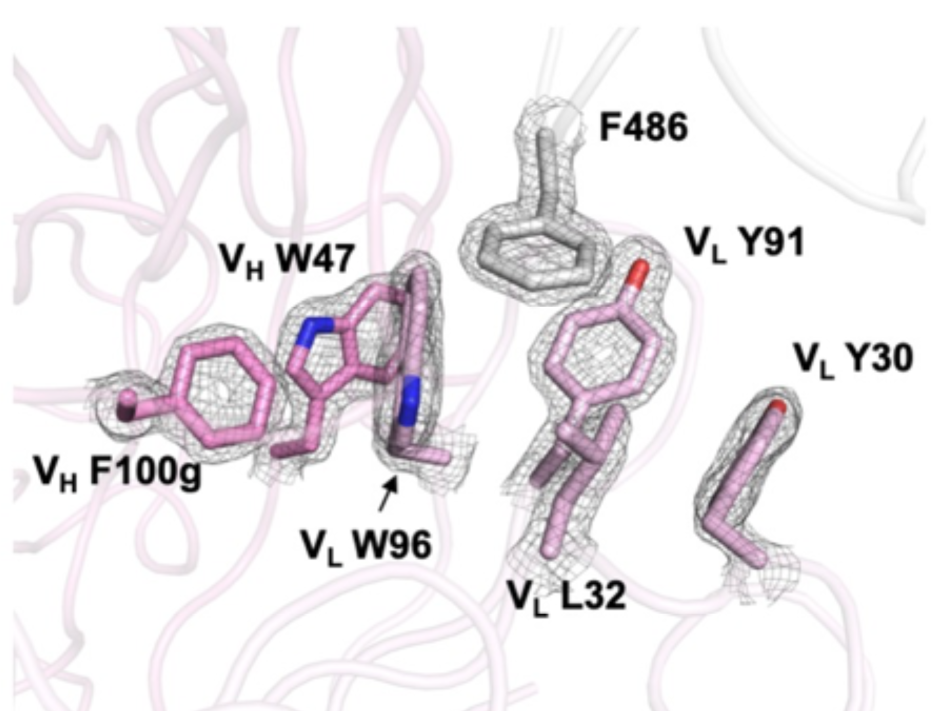
Electron density maps for the ridge-anchoring pocket. Final 2Fo-Fc electron density map for the ridge-anchoring pocket around RBD F486 is contoured at 2.0 σ.

**Supplementary Table 1.**
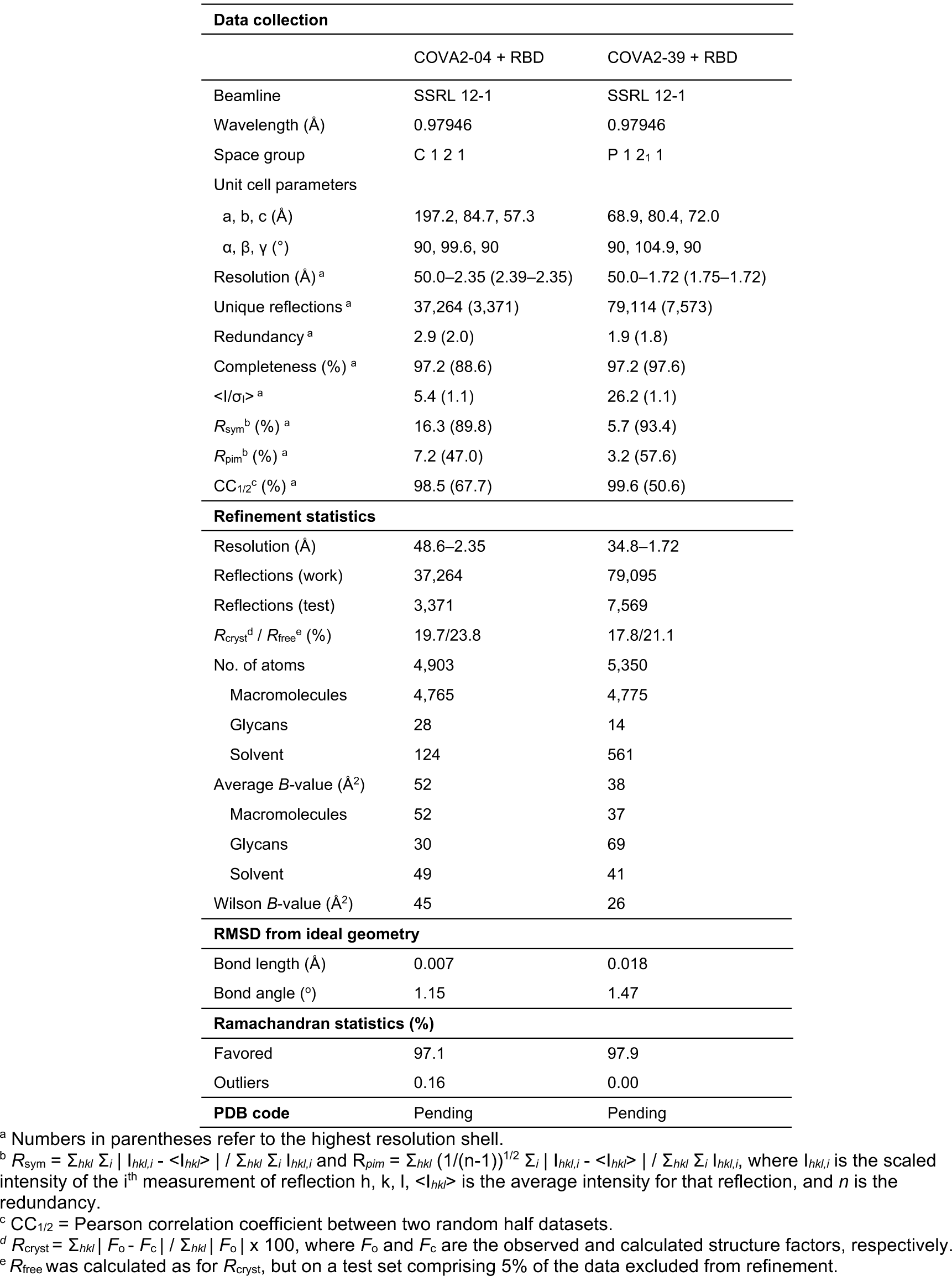
X-ray data collection and refinement statistics.

**Supplementary Table 2.** Hydrogen bonds and salt bridges identified at the antibody-RBD interface using the PISA program.

| COVA2-04 | Distance [Å] | SARS-CoV-2 RBD |
| --- | --- | --- |
| Hydrogen bonds |  |  |
| H:THR28[N] | 3.1 | A:ALA475[O] |
| H:ASN32[ND2] | 3.1 | A:ALA475[O] |
| H:TYR33[OH] | 2.6 | A:LEU455[O] |
| H:SER53[N] | 3.5 | A:TYR421[OH] |
| H:SER53[OG] | 3.1 | A:TYR421[OH] |
| H:SER53[OG] | 2.4 | A:ARG457[O] |
| H:GLY54[N] | 2.9 | A:TYR421[OH] |
| H:SER56[OG] | 2.3 | A:ASP420[OD2] |
| H:ARG94[NH1] | 3.1 | A:ASN487[OD1] |
| H:ARG94[NH2] | 3.2 | A:ASN487[OD1] |
| H:ARG94[NH2] | 3.7 | A:TYR489[OH] |
| H:ARG98[NE] | 3.9 | A:GLN493[OE1] |
| H:ARG98[NH1] | 3.1 | A:SER494[O] |
| H:GLY26[O] | 2.6 | A:ASN487[ND2] |
| H:SER31[O] | 2.7 | A:TYR473[OH] |
| H:GLU97[OE1] | 3.2 | A:LYS417[NZ] |
| H:GLU97[OE2] | 2.8 | A:TYR453[OH] |
| L:TYR32[OH] | 3.5 | A:TYR453[OH] |
| L:SER31[OG] | 3.6 | A:TYR495[O] |
| L:SER29[OG] | 3.8 | A:GLY496[O] |
| L:SER27A[OG] | 3.2 | A:THR500[O] |
| L:SER29[OG] | 3.7 | A:ASN501[OD1] |
| L:SER93[OG] | 2.7 | A:TYR505[OH] |
| L:SER93[N] | 3.8 | A:TYR505[OH] |
| L:GLY92[O] | 2.9 | A:ARG403[NH1] |
| L:SER93[OG] | 3.8 | A:ARG403[NH1] |
| Salt bridges |  |  |
| H:GLU97[OE1] | 3.2 | A:LYS417[NZ] |

| COVA2-39 | Distance [Å] | SARS-CoV-2 RBD |
| --- | --- | --- |
| Hydrogen bonds |  |  |
| H:THR53[N] | 3.4 | A:GLU484[OE2] |
| H:THR53[OG1] | 3.6 | A:GLU484[OE2] |
| H:GLY54[N] | 2.8 | A:GLU484[OE2] |
| H:THR56[N] | 3.3 | A:GLU484[OE1] |
| H:THR56[OG1] | 2.7 | A:GLU484[OE1] |
| H:TYR58[OH] | 2.6 | A:GLU484[O] |
| H:GLU100C[N] | 3.8 | A:ASN487[OD1] |
| H:GLU100C[N] | 2.9 | A:TYR489[OH] |
| H:GLU100C[OE2] | 3.0 | A:ASN487[N] |
| H:GLU100C[OE2] | 2.5 | A:TYR489[OH] |
| H:SER30[O] | 3.2 | A:GLN493[NE2] |
| L:SER95[OG] | 2.8 | A:PHE486[N] |

